# Evaluation of North American Asian Citrus Psyllid (Hemiptera: Liviidae) Population Genetics using cytochrome P450 melt curve analysis

**DOI:** 10.1101/445734

**Authors:** Juan F. Macias-Velasco, Ryan T. Brunson, Wayne B. Hunter, Blake R. Bextine

**Affiliations:** The University of Texas at Tyler, Department of Biology, Tyler TX; USDA- ARS, U.S. Horticultural Research Lab, 2001 South Rock Road, Fort Pierce, FL 34945, USA

**Author notes:** Corresponding author. Please address correspondence to: Blake Bextine, The University of Texas at Tyler, 3900 University Blvd, Tyler, TX 75799, Financial contact: Blake Bextine.

**Keywords:** Cytochrome P450, Asian Citrus Psyllid, HLB, Citrus Greening

## Abstract

The Asian Citrus Psyllid (ACP) (*Diaphorina citri*) is a Hemipteran which feeds on Citrus and is the principal vector of the pathogen *Candidatus* Liberibacter asiaticus (Las); the primary causal agent of Huanglongbing (HLB) or citrus greening disease. Currently HLB is the single greatest threat to the American citrus industry. Previous work has shown that that Cytochrome P450 (CYP4) genes are associated with insecticidal resistance to neonicotinoid insecticides. Infection by Las has also been shown to affect the expression of CYP4 genes. In this study the genetic relationships of ACP populations was evaluated using melting curve analysis of CYP4 genes. Principal component analysis shows the presence of two potential haplotypes. The groups identified by principal component analysis are significantly different from each other (P < 0.05) for all CYP4 genes tested. All populations are most likely within 2-3 mutations of each other. The existence of these mutations sheds light on the potential evolution of ACP in North America. Indicating either coevolution with Las or difference in imidacloprid basal selective pressure in the United States and Mexico.

---

The Asian Citrus Psyllid (ACP) (*Diaphorina citri*) is a Hemipteran that feeds on Citrus (Kuwayama, 1908). It is believed that ACP originated in Asia, but has spread to South America, Central America, North America, and the Middle East (Yang et al., 2006). It is also known to be the primary vector of the pathogen *Candidatus* Liberibacter asiaticus (Las); which is the causal agent of Huanglongbing (HLB) or Citrus Greening disease (Garnier & Bove, 1993; Jagoueix, Bove & Garnier, 1994). HLB is the most serious disease of citrus, first described in 1929 in China. It is characterized by a progressive decline of tree vigor, followed by inevitable death (Capoor, Rao & Viswanath, 1970). It has been estimated that HLB has resulted in over $3.63 billion in lost revenue (Hodges & Spreen, 2012) in the Florida citrus industry alone. As such HLB is currently the single greatest threat to the citrus industry. There is no cure for HLB. The great economic importance of this disease has led to the quarantine of citrus plant materials in Florida, Georgia, Puerto Rico, the U.S. Virgin Islands, two parishes in Louisiana, and two counties in South Carolina (Rosenthal & McNally, 2010). An additional quarantine has been put in place for ACP in the states of Alabama, Florida, Georgia, Hawaii, Louisiana, Mississippi, Texas, Guam, Puerto Rico and the U.S. Virgin Islands.

Given the current lack of methods to manage the pathogen, it is important to understand the movement and populations of the vector. Population genetics of ACP have previously been evaluated based upon the cytochrome oxidase subunit 1 (CO1) gene. This study concluded that though are at least two global populations of ACP (Boykin et al., 2012). They are from Southwest Asia (SWA) and Southeast Asia (SEA). North America is believed to have been invaded only by the SWA group and has limited genetic diversity.

Imidacloprid is a systemic insecticide within the class of chemicals known as neonicotinoids. It has previously been shown that Cytochrome P450 (CYP4) genes are associated with insecticidal resistance to neonicotinoids (Tiwari et al, 2011) in ACP. Infection by Las has also been shown to affect the expression of CYP4 genes (Tiwari et al, 2011). It would therefore be informative to evaluate the genetic relationships of ACP populations based upon said CYP4 genes. They are as follows: CYP4C68, CYP4DA1, CYP4DB1, and CYP4G70 (Accession #′s: JF934717, JF934718, JF934719, and JF934720).

One means by which genetic relationships can be evaluated is by ‘melt curve analysis’. This techniques is a well-established technique in the identification of genetic differences (Bextine & Child, 2007; Ramirez et al., 2010; Swisher, Munyaneza, & Crosslin, 2012; Tricarico et al., 2011) and has been previously used to identify potato psyllid (*Bactericera cockerelli*) haplotypes (Chapman, et al, 2012). In this study the genetic relationships of ACP populations was evaluated using melting curve analysis using the aforementioned CYP4 genes.

## Materials and Methods

Samples were collected from Los Angeles CA, San Bernardino CA, Riverside CA, Nuevo Leon MX, Bourg LA, Belle Chasse LA, New Orleans LA, and Horne á peau GP. Samples were stored in ethanol and subsequently stored at −80°C. Extraction and purification of DNA was done from individuals using a DNeasy Blood & Tissue Kit (Qiagen, Valencia CA). The DNA quality and quantity was determined with a nanodrop™ spectrophotometer (Thermo Scientific, Waltham MA).

Quantitative real-time PCR (qPCR) was performed using primers previously reported in (S Tiwari et al., 2011) for CYP4C68, CYP4DA1, CYP4DB1, and CYP4G70 (Accession #′s: JF934717, JF934718, JF934719, and JF934720). The reaction mixtures were composed of 12.5 μL Platinum^®^ SYBR^®^ Green qPCR SuperMix-UDG (Invitrogen™, Waltham MA), 2.5 μL (2.5 μM) of each primer, 6 μL sterile (DNASE, RNASE free) water (Fisher, Waltham MA), and 2.5 μL (∼25ng) of total genomic DNA. A Rotor-Gene Q (Qiagen, Valencia CA) real-time PCR thermal cycler was used with the following thermal profile: 50°C for 2 minutes, 95°C for 2 minutes, 40 cycles of 95°C for 15 seconds and 60°C for 30 seconds, followed by melt curve analysis in 0.5°C increments from 50-95°C. A preliminary qPCR was performed to confirm the identity of amplicons by melt curve analysis based on sequences reported previously. Gel electrophoresis was performed to further validate the amplicon identity by determining the molecular weight or the amplicons. Upon successful identification of amplicons, qPCR was performed on all subsequent samples and melting temperatures (T_m_) were determined.

Statistical significance between locations was determined by unpaired two-tailed student’s t-test. Principal component analysis (PCA) was performed using the R statistical software version 3.2.3 (R Core Team, 2013). PCA performed taking into account the T_m_ values alone of each sample in one scenario and T_m_, latitude, and longitude of each sample in a second scenario.

In order to determine the corresponding sequence differences resulting in differences in T_m_, the theoretical T_m_ of each CYP4 gene was determined by salt-adjusted methods reported in (Howley, Israel, Law, & Martin, 1979) assuming 50 mM salt concentration. Random G/A changes were made to the reference sequences (up to 5 changes) and the change in T_m_ resulting per G/A change (ΔG/A) was determined for each CYP4 amplicon by linear regression. The change in nucleotides (Δnt) relative to a respective reference was then calculated by subtracting the reference T_m_ from the sample T_m_ and dividing this difference by the respective ΔG/A.

## Results

Amplification was observed using total genomic DNA (Fig. 1A) and T_m_ values were observed ranging between 81.5°C and 83.5°C (Fig. 1C). As compared to the theoretical T_m_ values of the references sequences, which ranged between 80.2°C and 82.9°C. The molecular weight of CYP4C68, CYP4DA1, CYP4DB1, and CYP4G70 fell within the range of 100 to 200 base pairs (bp) (Fig. 1A)

**Fig. 1.**
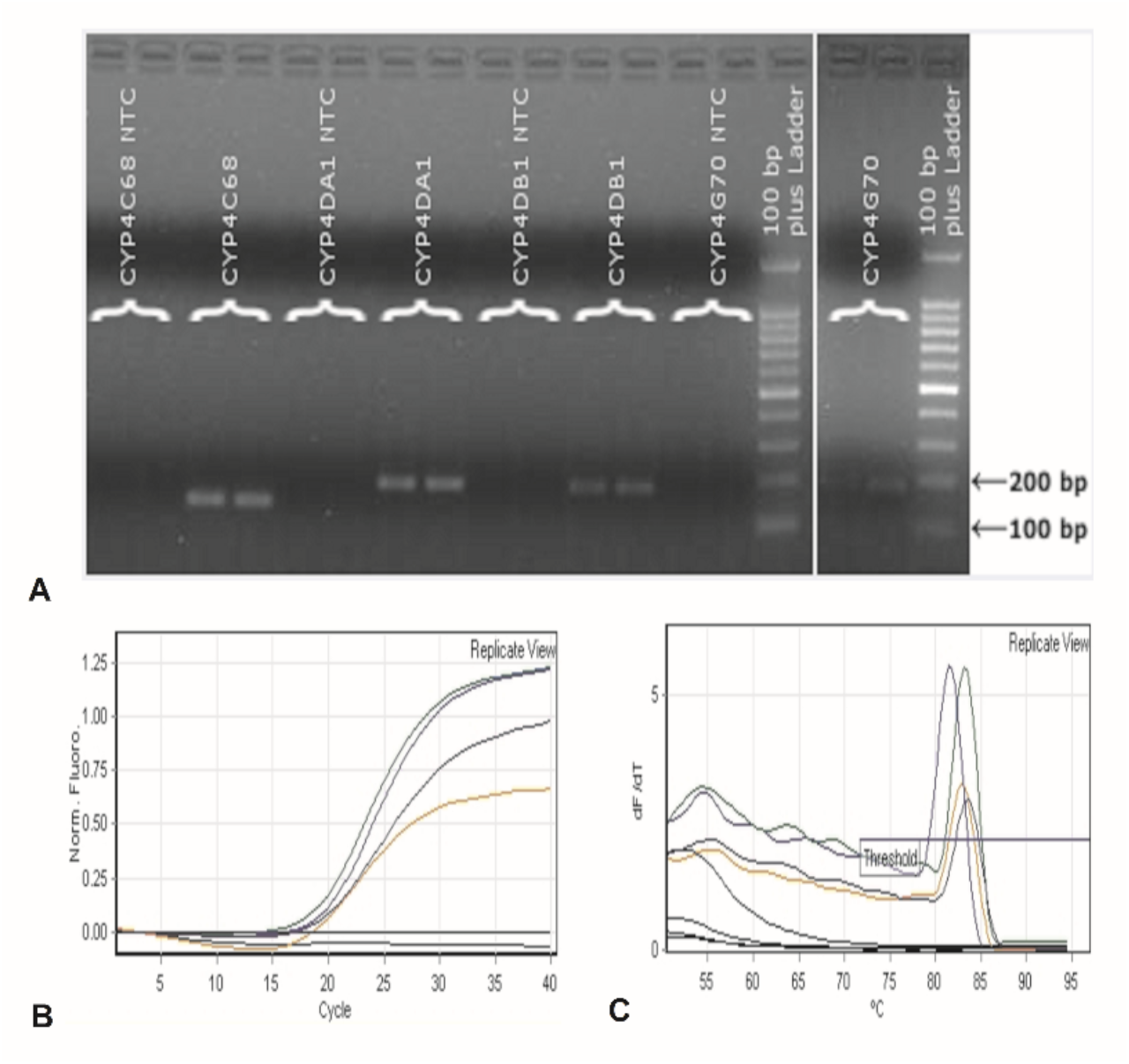
Confirmation of amplicon identity by melt curve analysis and molecular weight determination. Quantitative PCR was performed on total genomic DNA isolated from ACP samples. Quantitative PCR was performed for CYP4C68, CYP4DA1, CYP4DB1, and CYP4G70 using previously reported primers. Following the Quantitative PCR, melt curve analysis was performed from 50°C to 95°C in 0.5°C increments. Agarose gel electrophoresis was performed in order to determine the amplicon molecular weight. Identity was then assessed by comparing theoretical T_m_ and molecular weight to those observed. Clear amplification was observed for all four CYP4 genes (B). The observed T_m_ values were between 81.5°C and 83.5°C (C). As compared to the theoretical value range of 80.2°C and 82.9°C. The observed molecular weights fell within the range 100 to 200 bp (A). As compared to the theoretical value range of 152 bp to 194 bp. The amplicon molecular weights were all within 10 bp of the theoretical values (Data not shown).

The observed T_m_ values were 81.85-82.75°C, 80.60-81.35°C, 82.50-83.25°C, and 82.00-82.75°C (Fig. 2A.) for CYP4C68, CYP4DA1, CYP4DB1 and CYP4G70 respectively. The T_m_ values for San Bernardino were significantly different (P value < 0.0001) from other California samples for all four CYP4 genes. The T_m_ values for Nuevo Leon were significantly different (P value < 0.05) from Louisiana samples for all four CYP4 genes.

**Fig. 2.**
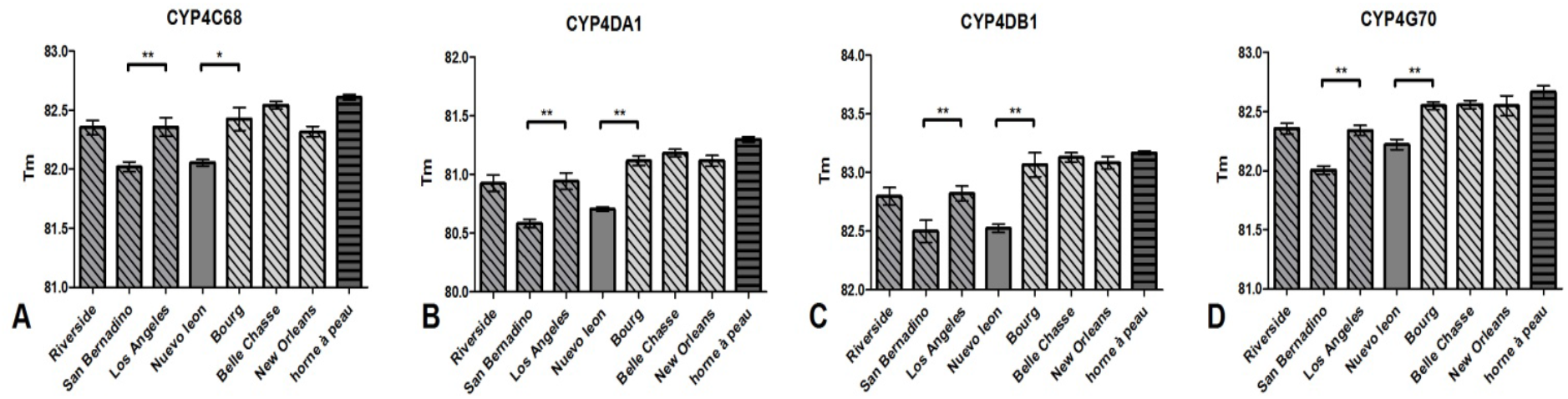
Determination of CYP4C68, CYP4DA1, CYP4DB1, and CYP4G70 T_m_ values by melt curve analysis. Quantitative PCR was performed on total genomic DNA isolated from ACP samples. Quantitative PCR was performed for CYP4C68, CYP4DA1, CYP4DB1, and CYP4G70 using previously reported primers. Following qPCR, melt curve analysis was performed from 50°C to 95°C in 0.5°C increments. Statistical significance between locations was determined by unpaired two-tailed student’s t-test. Determined T_m_ values were 81.85-82.75°C (A), 80.60-81.35°C (B), 82.50-83.25°C (C), and 82.00-82.75°C (D) corresponding to CYP4C68, CYP4DA1, CYP4DB1 and CYP4G70 respectively. Statistical significance noted by *, where ** is a P value < 0.0001 and * is a P value < 0.05.

Clustering by PCA, taking into account only T_m_ values, resulted in the observation of two distinct populations (Fig. 3A). Haplotype B, containing the majority of samples and Haplotype A, containing San Bernardino and Nuevo Leon samples. Clustering by PCA, taking into account latitude and altitude as well, yielded more defined clustering (Fig. 3B), but separation into haplotype A and B is still observed in the same manner. The proportions of each haplotype within the sampled regions show a clear geographical separation.

**Fig. 3.**
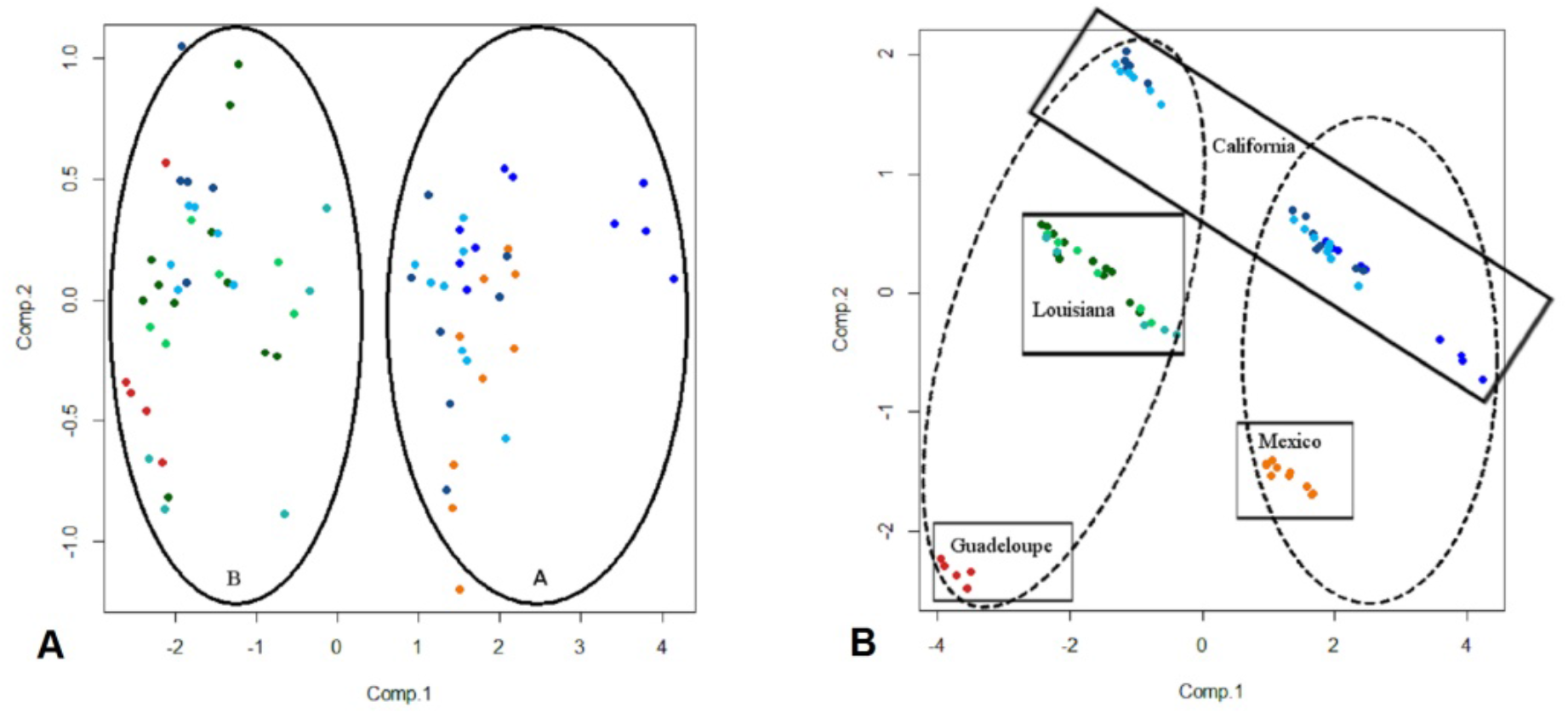
Clustering of ACP populations based on CYP4 genes, latitude, and longitude. Principal component analyses were performed using the T_m_ values of CYP4 genes without longitude and attitude (A) as factors, as well as with them as factors (B). Ellipsoids have no explicit statistical meaning. Additionally, Δnt values were extrapolated from T_m_ values to assess clustering based upon minimum sequence differences (C). Populations of ACP clustered into two groups regardless of the inclusion of latitude and longitude. Extrapolated minimum sequence differences demonstrate the same clustering pattern, wherein California and Mexico represent haplotype A, and all other sites represent haplotype B.

## Discussion

The original study from which primers were taken used complimentary DNA as the template, in this study genomic DNA was the template. The clear amplifications observed using these primer sets and the similarity to theoretical T_m_ values would indicate that intronic DNA is absent from these genes. This is further supported by the observed amplicon sizes, which were within ~14 bp of the predicted amplicon size (Figure 1). This would all show that the use of these previously reported primers with genomic DNA is a viable method to amplify CYP4 genes.

Principal component analysis shows the presence of two potential haplotypes (Figure 3 A&B) based on CYP4. The groups identified by principal component analysis are significantly different from each other for all CYP4 genes tested (P < 0.05) (Figure 2). The evolutionary distance is best understood in terms of how many base pair differences separate the two populations. To this end it was desirable to infer this distance by *in silico* random mutation and regression analysis. The result of extrapolating a minimum number of mutations between samples leads to the separation into haplotype A and haplotype B as expected. All populations are most likely within 2-3 mutations of each other. This limited diversity is in agreement with previous findings based on CO1. These mutations may shed light on the potential evolution of ACP in North America given that expression levels of these CYP4 genes have been shown to be associated with imidacloprid resistance in ACP (Killiny et al, 2014). The discovery of these haplotypes based upon these genes associated with imidacloprid resistance is indicative of potential differences of selection pressure. In particular it could be the result of different imidacloprid usage strategies. The implication being that Mexico has different strategies for the use of imidacloprid than the United States. This being the case, it could be inferred that populations may be moving from Mexico into southern California (Figure 4). Given that Las is known to affect expression of CYP4 genes, this may also be the result of co-evolution. Indicating that one population is more tolerant of Las infection than the other (Ammar et al, 2016; Arp et al, 2016; Mann et al, 2018). The exact effect of these changes and why this movement would not affect Louisiana populations is unknown. Alternatively, it may be the case that Haplotype A is a part of the SWE moving north from South America and Haplotype B is part of the SWA population moving west. Further analysis would need to be performed to correlate physiological and bacterial differences to these mutations.

**Fig. 4.**
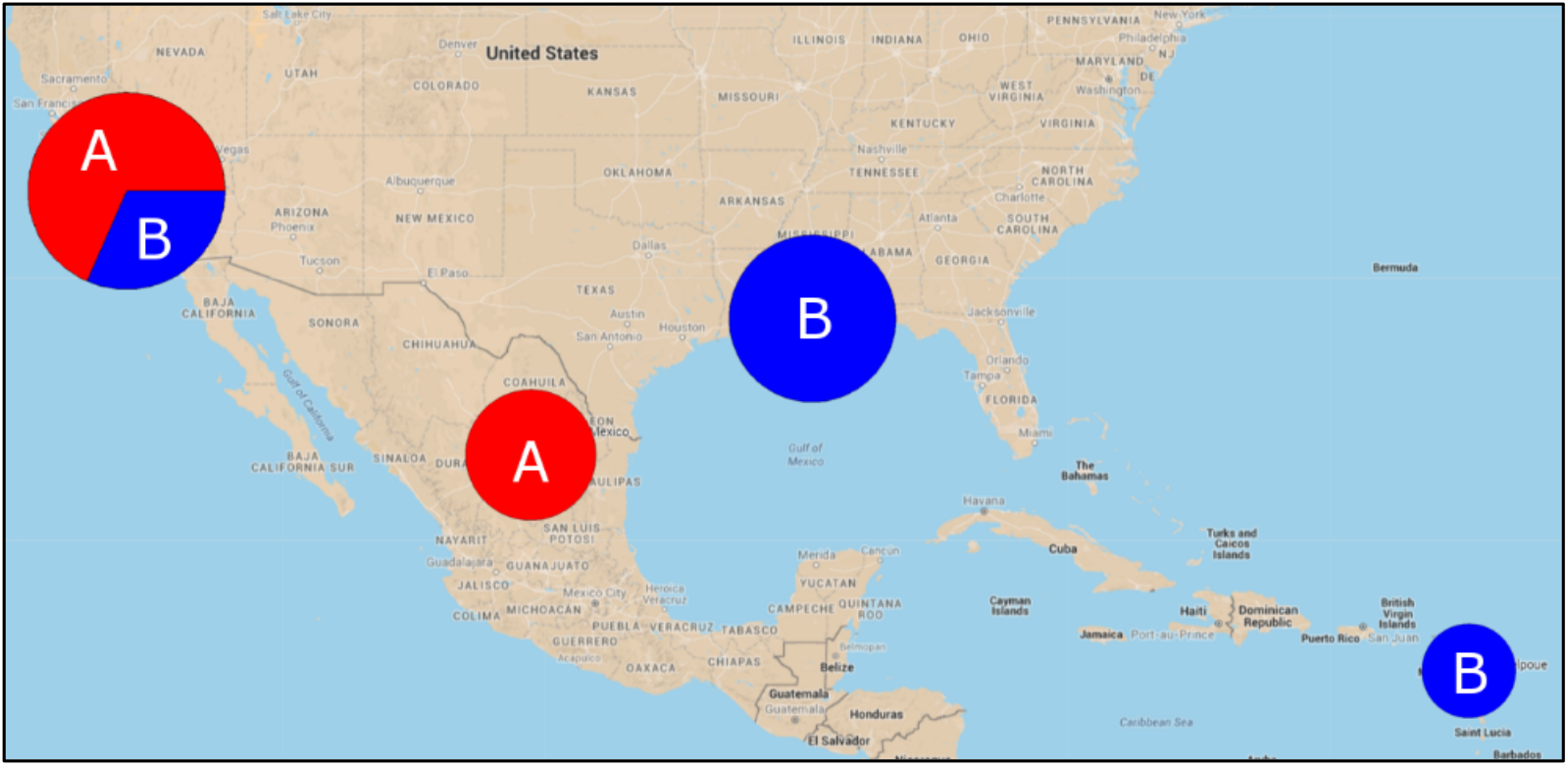
Composition of haplotypes amongst sampled regions. The number of each haplotype was determined based upon CYP4 genes T_m_ values as divided by PCA. Pie charts were generated for each sample region. The size of each pie chart was log_2_ scaled by sample size. Red and blue represent A and B haplotypes respectively.

## Acknowledgements

The authors would like to thank Chris Powell for providing guidance. This work was supported by USDA, ARS, U.S. Horticultural Research Laboratory, NP304_WH2046, Fort Pierce, FL.

